# COUNTERING AGE-ASSOCIATED ALTERATIONS IN OLIGODENDROCYTE-DERIVED EXTRACELLULAR MATRIX REJUVENATES COGNITION

**DOI:** 10.64898/2026.02.03.702938

**Authors:** Amber R. Philp, L. Remesal, Karishma J.B. Pratt, Jason C. Maynard, Shanan Sahota, Turan Aghayev, Gregor Bieri, Rebecca Chu, Gabriel Avillion, Juliana Sucharov-Costa, Yasuhiro Fuseya, Rhea Misra, Shir Mandelboum, Bende Zou, Julien Couthouis, Xinmin S. Xie, Stephen P.J Fancy, Alma L. Burlingame, Takakuni Maki, Saul A. Villeda

**Author notes:** Corresponding Authors: Dr. Saul Villeda, PhD, University of California San Francisco Department of Anatomy, 513 Parnassus Ave, Box 0452 San Francisco, CA 94143.

## Abstract

Efforts to rejuvenate age-related cognitive decline have predominantly targeted neurons, often overlooking non-neuronal cell types in the aging brain. Here, we show that countering alterations in oligodendrocyte-derived extracellular matrix (ECM) in the aging hippocampus restores cognition. We identify broad age-associated transcriptional and proteomic changes in oligodendrocytes, including dysregulation of the matrisome, with marked upregulation of ECM components and associated regulators with age. Among these, we detect an increase in Hyaluronan and proteoglycan link protein 2 (HAPLN2), an oligodendrocyte-derived core matrisome protein that locates specifically at the nodes of Ranvier, in the hippocampus of aged mice and older humans. Hapln2 overexpression in oligodendrocytes of young mice recapitulated age-related memory impairments. Conversely, abrogating the age-related increase in Hapln2 induced synaptic plasticity-related hippocampal transcriptional signatures and improved memory in aged mice. Together, these data define oligodendrocyte-derived ECM remodeling as a hallmark of brain aging that can be targeted to rescue cognitive decline.

## INTRODUCTION

Aging is the primary risk factor for cognitive decline^1,2^, yet the cellular and molecular mechanisms through which it impairs brain function with age remain poorly defined. Deciphering these mechanisms is crucial not only for understanding vulnerability to age-related neurodegenerative disease but also for identifying viable molecular targets to restore cognition in old age. Indeed, we and others have begun to challenge the longstanding view of brain aging as an inexorable and irreversible process by demonstrating that aspects of cognitive function and brain plasticity can be restored even in old age through systemic or cellular rejuvenation strategies^3–10^. However, these efforts to rejuvenate age-related cognitive function have been placed primarily on neurons. While neuronal dysfunction is undoubtedly central to brain aging, this emphasis has led to the relative underappreciation of non-neuronal cell types that contribute to functional decline in the aging brain.

Oligodendrocytes, traditionally viewed for their role in myelination, have received comparatively little attention in the context of cognitive aging. Beyond forming myelin, oligodendrocytes maintain axonal health and influence synaptic transmission efficiency through metabolic and structural support^11–16^. Emerging evidence reveals that oligodendrocytes undergo profound age-related changes that extend beyond myelin^17,18^, and that white matter tracts are focal hotspots of cellular aging^19^. However, most oligodendrocyte centered work has focused on white-matter regions, even though grey matter harbors abundant oligodendrocytes. The aging trajectories of these gray matter oligodendrocytes remain far less understood yet may contribute to age-related cognitive dysfunction through mechanisms distinct from demyelination.

Using single nucleus RNA sequencing (snRNA-seq), oligodendrocyte bulk RNA sequencing (RNA-seq), and mass spectrometry approaches, we identify dysregulation of the matrisome – the ensemble of extracellular matrix (ECM) and associated regulators – as a defining feature of aged oligodendrocytes. We observe an increase in the oligodendrocyte-derived core matrisome protein HAPLN2 specifically at the nodes of Ranvier in aged mice and older humans. Remarkably, modulating a single ECM component, Hapln2, broadly altered matrisome transcription and modulated cognition. Mimicking an age-related increase in oligodendrocyte-derived Hapln2 resulted in synaptic-related molecular changes and impaired cognitive performance in young mice. Excitingly, abrogating Hapln2 in aged oligodendrocytes induced synaptic plasticity-related transcriptional signatures and rescued age-related cognitive impairments. These findings indicate that alterations in oligodendrocyte-derived ECM are a critical driver of cognitive decline in aging.

## RESULTS

### Altered oligodendrocyte-derived extracellular matrix is a hallmark of hippocampal aging

To elucidate molecular mechanisms underlying oligodendrocyte aging that are associated with cognitive decline, we performed transcriptomic and proteomic analyses of the hippocampus – a brain region essential for learning and memory that undergoes profound age-related changes^20^. First, we performed snRNA-seq analysis on hippocampal nuclei from young (3 months) and aged (24 months) male mice (Figure 1A). A total of 14 cell clusters were identified using principal-component analysis (PCA)-based approach and projected by Uniform Manifold Approximation and Projection (UMAP) on a two-dimensional plot (Figure 1B), and populations were compared between conditions (Supplementary Figure 1A-E). Sub-clustering of the oligodendrocyte population revealed distinct age-related transcriptional signatures (Figure 1C), with differential expression analysis identifying widespread gene expression changes (Figure 1D). Gene Ontology (GO) enrichment analysis of upregulated and downregulated differentially expressed genes (DEGs) indicated that aging oligodendrocytes exhibited changes in cellular components associated with the plasma membrane and myelin sheath (Figure 1E). In contrast, the downregulated gene set was enriched for glutamatergic synapse, dendrite, and neuronal cell body (Figure 1F).

**Figure 1:**
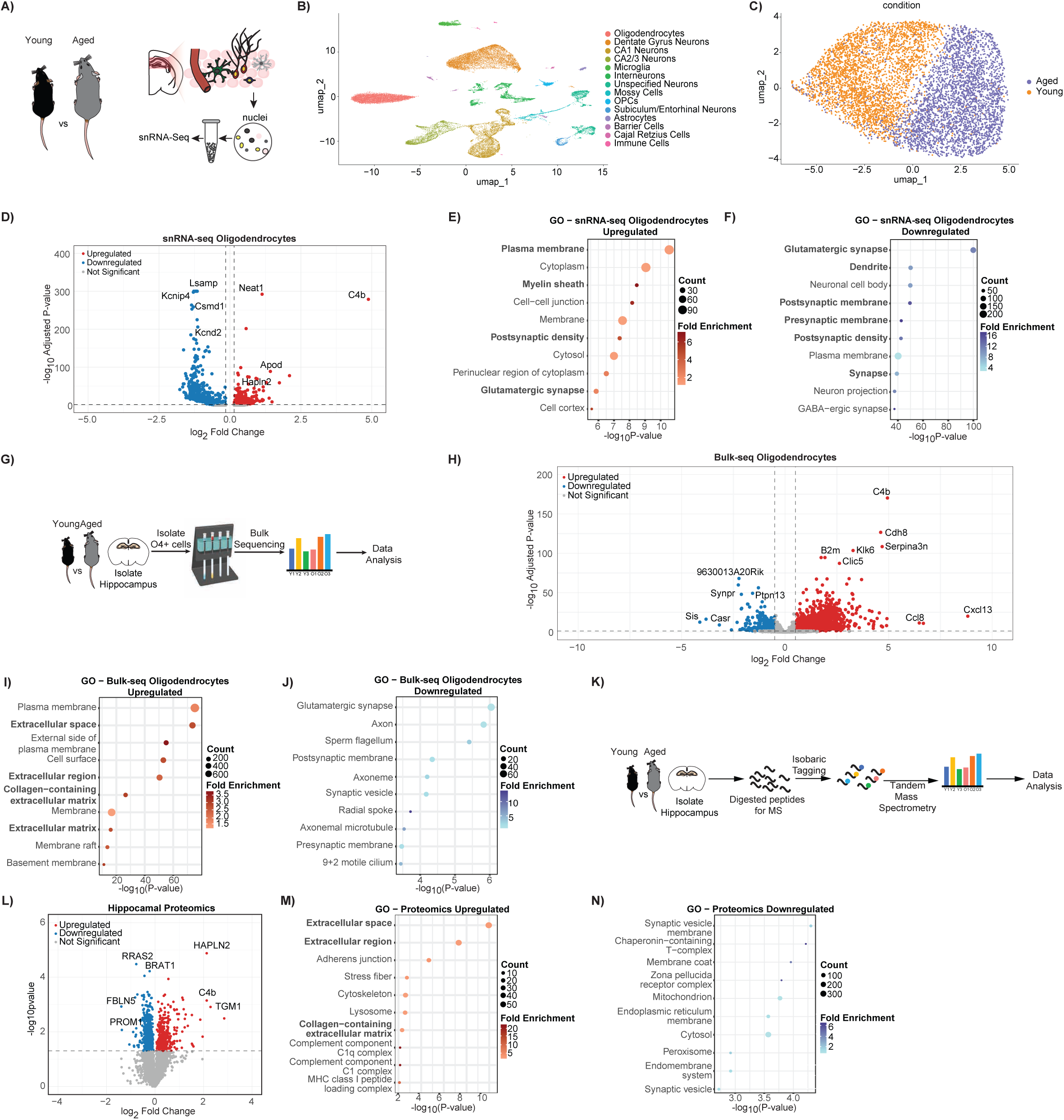
Multi-omics analysis identifies changes in oligodendrocyte-derived extracellular matrix as a hallmark of hippocampal aging. **A)** Schematic illustration of experimental groups, young (3-months) vs aged (24-months), and isolation of hippocampal nuclei for single-nucleus RNA sequencing (snRNA-seq). **B)** UMAP projection of identified hippocampal cell types. **C)** UMAP of subclustered oligodendrocytes separated by experimental group (young vs aged). **D)** Volcano plot of differentially expressed genes in oligodendrocytes identified from the snRNA-seq dataset shown in (B). **E–F)** Top Gene Ontology (GO) categories of cellular processes up- and down-regulated in oligodendrocytes with age based on snRNA-seq analysis. **G)** Overview of workflow for isolating hippocampal O4⁺ oligodendrocytes for bulk RNA sequencing. **H)** Volcano plot showing age-associated transcriptional changes in bulk-sequenced hippocampal O4⁺ oligodendrocytes. **I–J)** Top GO categories of cellular processes up- and down-regulated in bulk-sequenced hippocampal O4⁺ oligodendrocytes with age. **K)** Experimental workflow for quantitative proteomic analysis of hippocampal tissue from young and aged mice. **L)** Volcano plot showing age-associated changes in protein abundance in hippocampal tissue. **M–N)** Top GO categories of cellular processes up- and down-regulated in hippocampal proteins with age.

To extend these findings, we performed bulk RNA sequencing on oligodendrocytes isolated from the hippocampi of young and aged mice (Figure 1G). Assessment of magnetic-activated cell-sorted (MACS) O4-positive oligodendrocyte population confirmed enrichment for oligodendrocyte markers and depletion of transcripts associated with other major cell types in the hippocampus (Supplementary Figure 1F). Consistent with snRNA-seq data, bulk oligodendrocyte RNA-seq analysis revealed robust transcriptional differences with age (Figure IH), with cellular components associated with downregulated DEGs closely mirroring that of the snRNA-seq data (Figure 1J). However, we detected enrichment for extracellular matrix (ECM) related components among upregulated DEGs in the bulk sequenced oligodendrocyte population (Figure 1I).

Due to technical limitations in the number of animals necessary to obtain sufficient material for single-cell proteomic analysis of hippocampal oligodendrocytes, we instead performed quantitative proteomic profiling of soluble proteins from whole hippocampal tissue. This approach revealed widespread age-related changes in protein abundance (Figure 1K,L). Go analysis showed that downregulated proteins were primarily associated with synaptic signaling. In contrast, upregulated proteins were strongly enriched for ECM components, highlighting broad ECM remodeling as a defining molecular feature of the aged hippocampus (Figure 1M,N). Together, these integrated multi-omics analyses demonstrate that hippocampal aging is accompanied by coordinated transcriptional and proteomic remodeling of oligodendrocytes and their extracellular environment.

### Oligodendrocyte and hippocampal aging drive matrisome remodeling

Given the prominent upregulation of ECM genes and proteins in aged oligodendrocytes and hippocampus, we next sought to characterize age-associated changes within this molecular network. For this, we leveraged MatrisomeR^21^, a bioinformatic framework that systematically annotates ECM and ECM-associated molecules (collectively referred to as the matrisome). The matrisome comprises two broad categories: the core matrisome, which includes glycoproteins, collagens, and proteoglycans, and matrisome-associated components, encompassing ECM-affiliated proteins, ECM regulators, and secreted factors^22^ .

Leveraging our oligodendrocyte bulk RNA-seq dataset, we examined the matrisome composition of oligodendrocytes (Figure 2A). Next, we focused on genes associated with the extracellular space by GO analysis and detected broad transcriptional shifts between young and aged oligodendrocytes (Figure 2B). Differential expression analysis using MartisomeR further identified numerous matrisome transcripts that were significantly altered with age (Figure 2C). When we classified and normalized the number of differentially expressed matrisome genes to the total number of detected genes in each category (Supplementary Figure 2A), we observed widespread transcriptional upregulation across both core matrisome and matrisome-associated components in aged oligodendrocytes (Figure 2D,E). To determine whether transcriptional changes in aged oligodendrocytes were reflected at the protein level in the aged hippocampus, we examined the hippocampal matrisome proteome (Figure 2F) and observed that its distribution broadly mirrored the oligodendrocyte matrisome, with glycoproteins and ECM regulators again representing major components (Figure 2A,F). Proteins annotated to the extracellular space by GO analysis displayed pronounced age-related differences (Figure 2G), with differential expression analysis revealing strong upregulation of matrisome proteins (Figure 2H). When assessing the proportional distribution of altered ECM components in the aged hippocampus, relative to the total number of detected matrisome proteins (Supplementary 2B), we again observed an overall upregulation in both core matrisome and matrisome-associated components, most prominently within the proteoglycan and ECM-regulator classes (Figure 2I). Together, these multi-omics data indicate that aging is accompanied by broad transcriptional and proteomic expansion of the oligodendrocyte-derived and hippocampal matrisome, reflecting widespread age-related remodeling of ECM architecture and signaling pathways.

**Figure 2:**
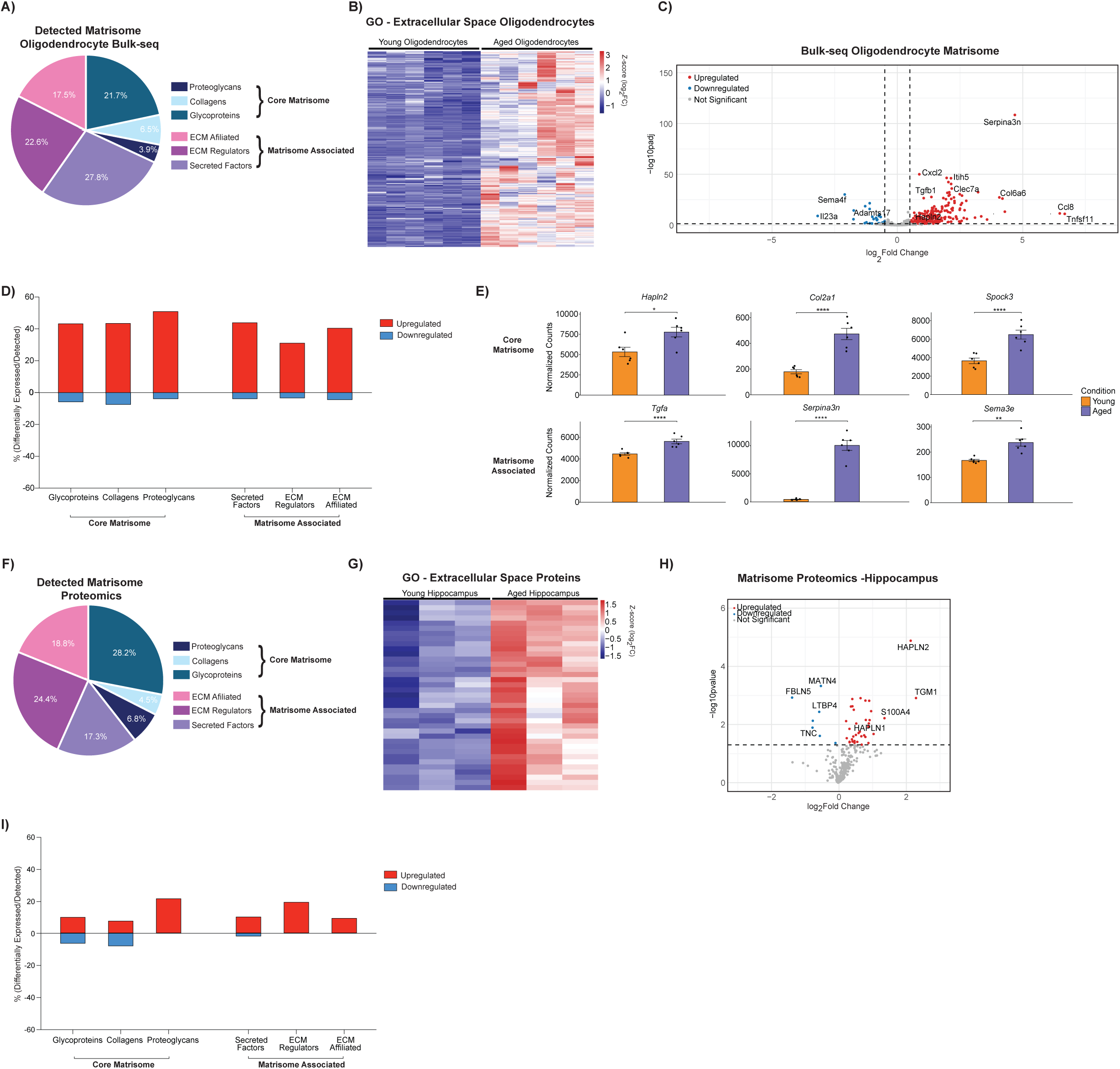
Transcriptomic and proteomic analysis reveal age-associated remodeling of the hippocampal matrisome. **A)** Pie chart showing the composition of matrisome genes detected in oligodendrocyte bulk RNA-seq. **B)** Heatmap of genes belonging to the *Extracellular Space* Gene ontology (GO) term from bulk-sequenced hippocampal O4⁺ oligodendrocytes. **C)** Volcano plot of differentially expressed Matrisome genes in bulk-sequenced hippocampal O4⁺ oligodendrocytes with age. **D)** Percentage of up- and down- regulated Matrisome genes with aging in bulk-sequenced O4⁺ oligodendrocytes, normalized to the total number of detected genes within each Matrisome category. **E)** Bar plots showing representative Matrisome genes upregulated with aging in bulk-sequenced O4⁺ oligodendrocytes. Differential expression significance was determined by DESeq2 using adjusted p-values. **F)** Pie chart showing the composition of matrisome proteins detected in hippocampal tissue. **G)** Heatmap of proteins identified in the extracellular space GO term from proteomic analysis of hippocampal tissue. **H)** Volcano plot of differentially expressed Matrisome proteins in hippocampal tissue with age. **I)** Percentage of up-and down- regulated Matrisome proteins with aging in hippocampal tissue, normalized to the total number of detected proteins within each category. Data are shown as mean +/- s.e.m. * p < 0.05, ** p < 0.01, *** p < 0.001, **** p < 0.0001.

### Increased oligodendrocyte-derived nodal ECM protein HAPLN2 alters hippocampal transcriptional profiles and impairs cognition

Having established that aging is accompanied by broad remodeling of oligodendrocyte-derived and hippocampal ECM, we next sought to identify potential pro-aging factors. We reasoned that molecules consistently altered across our oligodendrocyte snRNA-seq, bulk oligodendrocyte RNA-seq, and hippocampal proteomic datasets could represent core mediators of age-related hippocampal dysfunction. Comparative analysis identified one factor with conserved age-associated upregulation: the oligodendrocyte-derived core matrisome protein HAPLN2 (Figure 3A). HAPLN2 stabilizes ECM structures surrounding the nodes of Ranvier and is known to play an essential role in maintaining neuronal conductivity^23^. Using immunofluorescence (IF) staining we further confirmed the presence of HAPLN2 at the perinodal space (Supplementary 3A). Consistent with our multi-omics analysis, we observed increased Hapln2 expression in the hippocampus with age by Western blot analysis, a pattern that was also evident in aged female mice (Figure 3B,C, Supplementary Figure 3B-E). To gain further regional resolution, we performed immunohistochemical analysis in the hippocampus of young and aged mice (Figure 3D,E). We observed an age-related increase in Hapln2, specifically at the nodes of Ranvier, across hippocampal subregions (DG, CA1, CA2 and CA3), with the most pronounced increase detected in the CA2 and CA3 (Figure 3D,E). Next, to investigate the conservation of our findings in humans, we assessed changes in nodal HAPLN2 in the hippocampus of young and older individuals and similarly detected an age-related increase across hippocampal subregions (Figure 3F,G).

**Figure 3:**
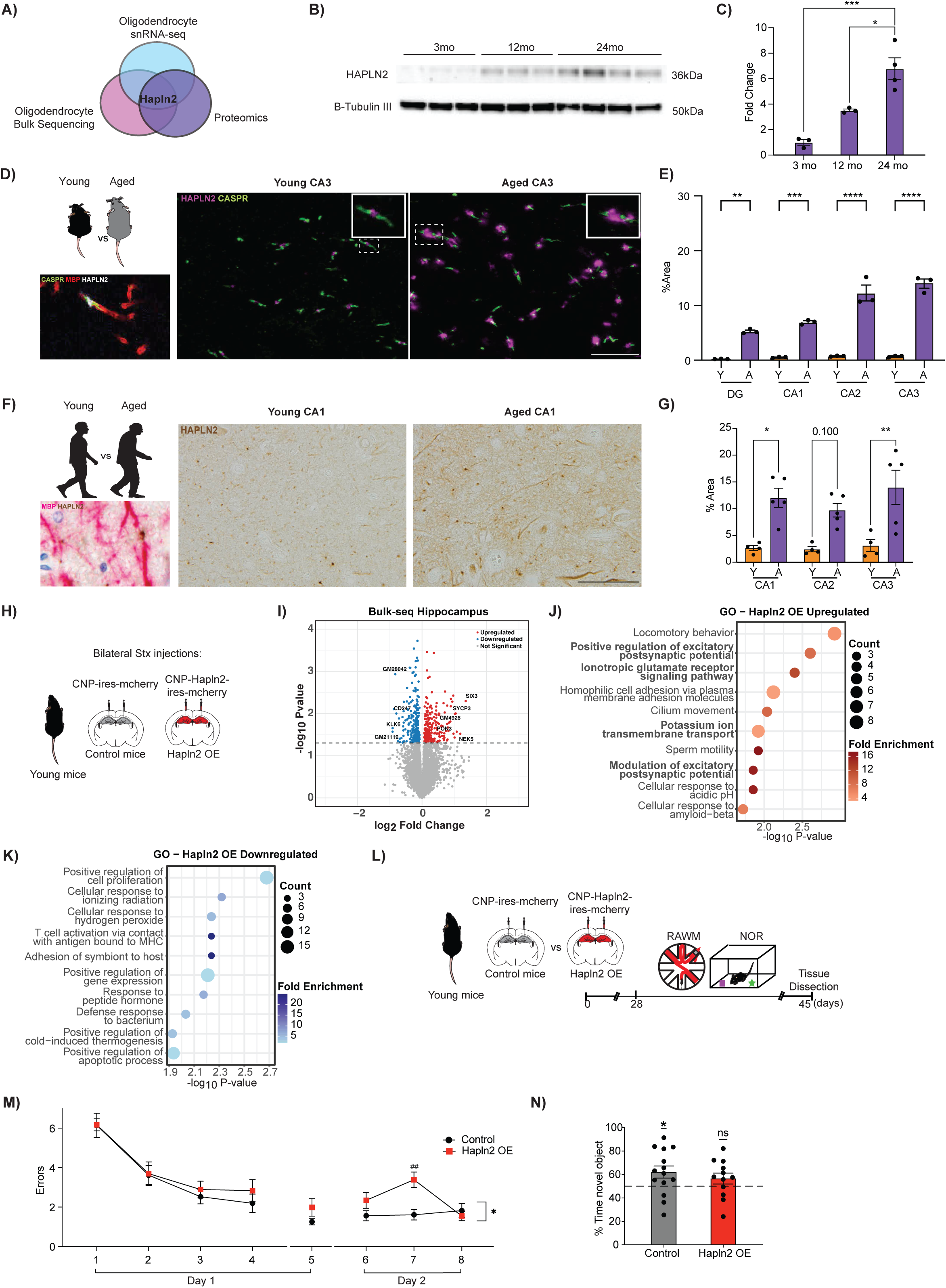
Mimicking an age-related increase in the core matrisome protein Hapln2 in oligodendroctyes impairs cognition in young mice. **A)** Hapln2 expression increases with age across oligodendrocyte snRNA-seq, O4⁺ oligodendrocyte bulk sequencing, and hippocampal proteomics. **B)** Western blot analysis of HAPLN2 protein levels in hippocampal tissue across age. **C)** Quantification of Western Blot shown in (B). **D)** Representative IF images showing HAPLN2 localization at nodes of ranvier in mice, delineated by MBP (myelin sheath) and CASPR (paranodes) in the hippocampal CA3 region (left). Representative HAPLN2 staining at the node of ranvier in young and aged CA3 hippocampal sections (right). Scale bar 10 μm. **E)** Quantification of changes in HAPLN2 across mouse hippocampal subregions with age. **F)** Representative immunohistochemistry images show HAPLN2 localization at nodes of ranvier in humans, delineated by MBP, in the CA1 region (left). Representative HAPLN2 staining in human CA1 (right). Images correspond to a 34-year-old male and a 95-year-old male. Scale bar 50 μm. **G)** Quantification of changes in HAPLN2 across human hippocampal subregions with age. **H)** Experimental design for lentivirus overexpression of *Hapln2* under the oligodendrocyte-specific *CNP* promoter in the hippocampus of young mice (2-3 mo). **I)** Volcano plot of differentially expressed genes in the hippocampus of Hapln2 overexpression (OE) mice compared to control mice. **J-K)** Top GO categories of biological processes associated with up- and down-regulated genes in Hapln2 OE mice. **L)** Experimental timeline of hippocampal stereotaxic injections and subsequent cognitive testing of young Hapln2 OE and control mice. **M)** Hippocampal-dependent spatial memory and learning was evaluated by radial arm water maze as number of errors while locating the escape platform. **N)** Object recognition memory was assessed by NOR. N= 12-14 mice. Data shown as mean +/- s.e.m. Statistical analysis was performed using one-way ANOVA with Tukey’s multiple comparisons (C, E, G), two-way ANOVA with Šidák’s post hoc test (M), or one-sample t-test vs 50% (M). * p < 0.05, ** p < 0.01, *** p < 0.001, **** p < 0.0001.

We next investigated whether Hapln2 overexpression in oligodendrocytes of young mice was sufficient to induce aging-like phenotypes in the hippocampus using a cell type specific, viral-mediated overexpression approach. Young mice were given bilateral stereotaxic injections of lentivirus encoding *Hapln2* or *mCherry* under the oligodendrocyte-specific 2′,3′-cyclic nucleotide 3′-phosphodiesterase (CNP) promoter into the hippocampus and molecular, and cognitive assessment was performed one month post injection (Figure 3H). Hapln2 overexpression was validated *in vivo* (Supplementary Figure 4A-C). For molecular analysis, we performed bulk RNA-seq analysis of hippocampal tissue and detected transcriptional alterations in Hapln2-overexpressing (OE) mice compared to control animals (Figure 3I). GO analysis of DEGs indicated enrichment for pathways involved in neuronal excitability and synaptic signaling, including *positive regulation of excitatory postsynaptic potential*, *potassium ion transmembrane transport*, and *modulation of excitatory postsynaptic potential* (Figure 3J,K). These findings suggest that Hapln2 OE alters neuronal function within the hippocampus. Functionally, we assessed whether Hapln2 overexpression in young mice affected hippocampal-dependent learning and memory using radial arm water maze (RAWM) and novel object recognition (NOR) behavioral paradigms (Figure 3L). Young control animals exhibited normal hippocampal-dependent learning and memory (Figure 3M,N). However, Hapln2 OE mice committed more errors when locating a hidden platform during RAWM testing (Figure 3M) and showed a decreased bias for the novel object in NOR task (Figure 3N), indicating deficits in spatial and object memory. No differences in overall wellbeing metrics were observed between groups (Supplementary 5A-E). Together, these findings indicate that mimicking an age-related increase in oligodendrocyte Hapln2 recapitulates in part molecular and cognitive features associated with hippocampal aging in young mice.

### Targeting Hapln2 in oligodendrocytes modulates the hippocampal matrisome, elicits synaptic-related transcriptional profiles and improves cognition at old age

Conversely, we investigated the possibility of rescuing aging phenotypes in the aged hippocampus by reducing Hapln2 expression in aging using an adeno-associated virus (AAV)-mediated shRNA approach. Hapln2 knockdown (KD) was validated *in vitro* and *in vivo* (Supplementary Figure 6A-E). Aged (22 months) mice were given retro-orbital injections of AAV-PHP.eB encoding shRNA sequences targeting Hapln2 or scramble control sequences. (Figure 4A). Bulk RNA-seq analysis was performed on hippocampal tissue and widespread transcriptional changes were observed in Hapln2 KD aged mice compared to controls (Figure 4B), with GO analysis of downregulated DEGs identifying ECM organization as one of the top associated biological processes (Figure 4C-D). Next, we characterized matrisome gene representation in the hippocampal bulk RNA-seq dataset (Figure 4E). When we assessed the proportional distribution of altered matrisome genes following Hapln2 abrogation, normalized to the total number detected for each category (Supplementary Figure 2C), we observed a pronounced and coordinated reduction in expression of matrisome genes spanning core matrisome and matrisome-associated components (Figure 4F,G). In addition to ECM-related pathways, GO analysis of downregulated DEGS identified processes associated with immune activity. However, immunohistochemical analysis of the aged hippocampus revealed no changes in microglial activation (IBA1) or complement C1q expression between Hapln2 KD and control animals (Supplementary Figure 7A,B).

**Figure 4:**
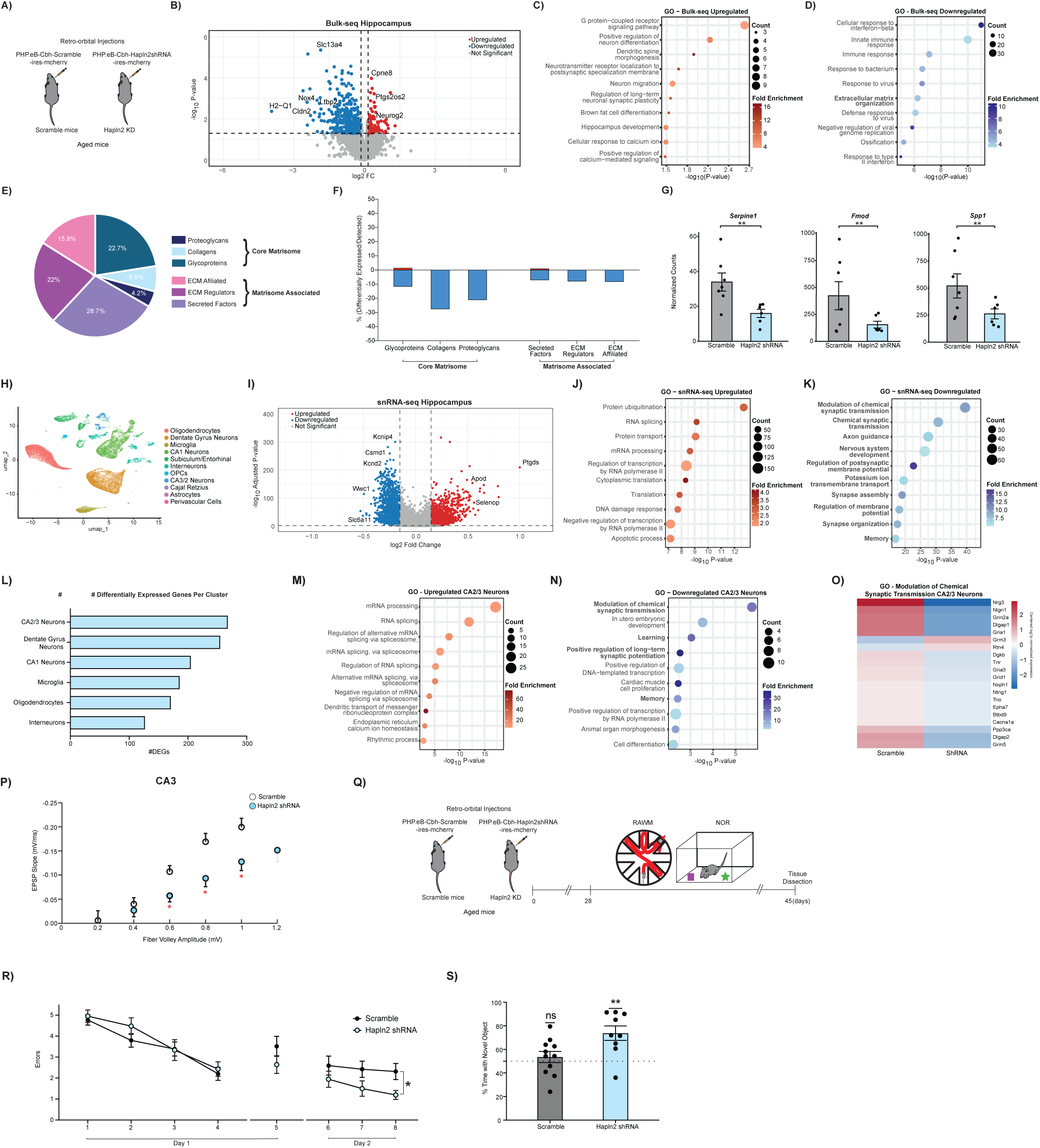
Targeting Hapln2 in aged mice remodels hippocampal ECM and rescues age-related cognitive impairments. **A)** Experimental schematic of Hapln2 knockdown in aged mice (22-24 mo) using AAV-PHP.eB carrying a Hapln2 shRNA or scramble control. **B)** Volcano plot of differentially expressed genes from bulk hippocampal RNA sequencing of Hapln2 shRNA mice compared to scramble treated mice. **C-D)** Top GO categories of biological processes associated with up- and down-regulated genes in Hapln2 shRNA treated mice. **E)** Pie chart showing the composition of matrisome genes detected in hippocampal bulk-seq. **F)** Percentage of up- and down- regulated matrisome genes in Hapln2 shRNA treated hippocampi, normalized to the total number of detected genes within each matrisome category. **G)** Bar plots showing representative matrisome genes downregulated in Hapln2 shRNA hippocampi. Differential expression significance was determined by DESeq2 using raw p-values. **H)** UMAP projection of identified hippocampal cell types in Hapln2 shRNA and scramble treated mice. **I)** Volcano plot of differentially expressed genes identified by snRNA-seq across hippocampal cell clusters following Hapln2 knockdown. **J-K)** Top GO categories of biological processes associated with up- and down-regulated detected by snRNA-seq in Hapln2 shRNA treated mice. **L)** Number of differentially expressed genes per cluster following Hapln2 knockdown. **M-N)** Top GO categories of biological processes associated with up- and down-regulated genes in the CA2/3 neuronal cluster. **O)** Heatmap of genes belonging to the *modulation of chemical synaptic transmission* GO term in CA2/3 neurons. **P)** Input–output relationship of EPSP slope plotted against fiber volley amplitude recorded in the CA3 region. **Q)** Experimental timeline of retro-orbital injections and subsequent cognitive testing of aged Hapln2 shRNA and scramble mice. **R**) Hippocampal-dependent spatial memory and learning was evaluated by radial arm water maze as number of errors while locating the escape platform. **S)** Object recognition memory was assessed by NOR. N= 9-11 mice. Data shown as mean +/- s.e.m. Statistical analysis was performed using two-way ANOVA with Šidák’s post hoc test (L) or one-sample t-test vs 50% (M). * p < 0.05, ** p < 0.01.

To further characterize molecular changes at a cell type specific level, we performed snRNA-seq analysis of the aged hippocampus following abrogation of Hapln2. Analysis identified a total of 11 cell clusters corresponding to major hippocampal cell types and populations were compared between treatment groups (Figure 4H, Supplementary 8A-E). Differential expression analysis across all cell types revealed widespread transcriptional alteration in Hapln2 KD compared to control mice (Figure 4I). GO analysis of downregulated DEGs identified prominent changes in neuronal processes, including *modulation of chemical synaptic transmission*, *axon guidance*, and *regulation of postsynaptic membrane potential* (Figure 4J,K). We next compared the number of DEGs across cell types and observed that the most extensive transcriptional changes occurred within excitatory neuron populations, with the CA2/3 neuronal cluster showing the highest number of DEGs (Figure 4L). GO analysis of downregulated DEGs specific to the CA2/3 excitatory neuron population again highlighted biological processes related to *synaptic transmission*, *learning*, and memory (Figure 4M,N). To further resolve these changes, we visualized the top 20 genes contributing to the GO term “*modulation of chemical synaptic transmission*” (Figure 4O). The resulting heatmap revealed a coordinated downregulation of genes mediating synaptic signaling in Hapln2 KD mice, consistent with impaired excitatory transmission within the CA2/3 circuit.

To complement transcriptional analysis, we performed field excitatory postsynaptic potential (fEPSP) recordings in acute hippocampal slices. Consistent with molecular changes in CA2/3 neuronal populations, Hapln2 KD mice exhibited reduced EPSP slope relative to fiber volley amplitude in the CA3 region, suggesting that larger action potentials is needed to propagate CA3 excitation (Figure 4P–O).Given that age-related increased excitability in the CA3 region is linked to cognitive deficits^24,25^ we asked whether attenuation of this hyperexcitability following Hapln2 abrogation could result in cognitive improvements. Hippocampal-dependent learning and memory was assessed in aged Hapln2 KD and control mice by RAWM and NOR testing (Figure 4Q). In the training phase of the RAWM paradigm, all mice showed similar spatial learning ability (Figure 4R). However, while aged control mice exhibited cognitive impairments, aged Hapln2 KD mice demonstrated improved learning and memory for the platform location committing less errors during the testing phase of the task compared with control animals (Figure 4R). Similarly, during NOR testing, aged control mice demonstrated impaired object memory showing no object preference, while aged Hapln2 KD mice were biased towards a novel object relative to a familiar object (Figure 4S). No differences in overall wellbeing metrics were observed between groups (Supplementary Figure 9A-E). Together, these data indicate that targeting the age-related increase in the oligodendrocyte-derived ECM protein Hapln2 can rescue molecular, functional, and cognitive hippocampal deficits at old age.

## DISCUSSION

Cumulatively, our data identify age-associated remodeling of the oligodendrocyte-derived ECM as a driver of hippocampal cognitive decline. Moreover, these data posit the nodal core matrisome protein HAPLN2 as an attractive molecular target by which to reverse memory loss late in life.

Cognitive decline in aging is linked to age-related neuronal dysfunction at the synaptic level^20,26^. As such, focus has been predominantly placed on identifying molecular mechanisms in neurons that can be targeted to restore cognition. Indeed, we and others have begun to identify factors, such as CREB, GNAQ, OGT, and FTL1 whose neuron-specific manipulation can elicit cognitive improvements^5,20,27–29^. While neuronal dysfunction is undoubtably a critical factor in brain aging, this emphasis has led to a relative underappreciation of non-neuronal cell types that contribute to age-related functional decline and could provide yet unexplored cellular and molecular targets to combat cognitive impairments. In this context, our work identifies oligodendrocytes as critical players in driving cognitive decline upstream of neuronal alterations through the modulation of their ECM and subsequent disruption of synaptic plasticity.

The ECM in the adult brain is essential for proper regulation of synaptic plasticity, neuronal signaling, and memory^30–34^, with age-related ECM remodeling poised to exert profound effects on cognitive function. Aging is accompanied by progressive accumulation and crosslinking of ECM proteins, leading to increased tissue stiffness and reduced molecular diffusivity, features that constrain neural plasticity^35–38^, remyelination, lymphatic drainage, and angiogenesis in the brain^38–41^. Much of our current understanding on ECM remodeling in the aging brain stems from work on microglia^42^. In the hippocampus, microglia promote synaptic remodeling through activity dependent degradation of perisynaptic ECM components such as chondroitin sulfate proteoglycans, a process essential for maintaining structural plasticity. However, during aging, diminished microglia capacity to remodel synaptic ECM results in excessive stabilization and reduced synaptic plasticity, contributing to memory consolidation^43,44^. In the cerebellum, elevated hyaluronic acid levels and age-related HAPLN2 aggregation induced microglia inflammatory responses^45^. Of note, in our study, cognitive rejuvenation by targeting HAPLN2 in hippocampal oligodendrocytes was not accompanied by changes in microglia activation markers, suggesting region specific regulation of downstream synaptic and inflammatory processes, and underscoring the need to further investigate age-related changes in oligodendrocyte ECM interactions.

Oligodendrocytes and their precursors synthesize key ECM constituents -including collagens, glycoproteins, and link proteins -that assemble the perinodal matrix surrounding nodes of Ranvier ^46,47^. Notably, manipulation of the transcription factor, Sp7, which regulates nodal ECM gene expression, was recently shown to alter ECM composition and brain stiffness, demonstrating that even highly localized ECM changes at nodes of Ranvier can exert widespread biophysical effects on neural tissue^48^. Consistent with this concept, our findings show that Hapln2 abrogation in aged mice triggers extensive remodeling of core matrisome and matrisome-associated components in the hippocampus, spanning multiple ECM classes and their regulatory factors. These results highlight that oligodendrocyte-derived ECM remodeling, though anatomically focal, can induce broad structural and molecular adaptations in the aging brain that have robust functional consequences on cognition.

From a translational perspective, aging is the most dominant risk factor for age-related neurodegenerative disorders^49^. Beyond its role in physiological aging, HAPLN2 dysregulation has been implicated in neurodegenerative disease, indicating that ECM remodeling may represent a convergent mechanism between aging and pathology. Elevated HAPLN2 expression has been reported in Parkinson’s disease, and HAPLN2 upregulation in mouse models of Parkinson’s disease pathology promotes α-synuclein aggregation and dopaminergic neuron vulnerability^50,51^. Thus, our data highlight the possibility of targeting oligodendrocyte-derived ECM components as a new therapeutic avenue for both restoring age-related cognitive decline and ameliorating neurodegenerative disease pathology in older individuals.

## RESOURCE AVAILABILITY

### LEAD CONTACT

Further information and requests for resources and reagents should be directed to and will be fulfilled by the lead contact, Dr. Saul Villeda.

## MATERIALS AVAILABILITY

Plasmids generated in this study are available from the lead contact upon request.

## DATA AND CODE AVAILABILITY

- The single-nucleus and single-cell RNA-sequencing datasets are available at the Gene Expression Omnibus (GEO). Accession numbers are listed in the <u>key resources table</u>.
- This paper does not report original code.
- Additional details are available from the <u>lead contact</u> upon request.

## ACKNOWLEDGMENTS

We thank Dr. Valerie Weaver for critical input on ECM analyses. Work was funded by Simons Foundation (S.A.V.), Bakar Family Foundation (S.A.V.), Hevolution Foundation (S.A.V.), Multiple Sclerosis Foundation (A.P), Frontiers in Medical Research fellowship (K.J.B.P), Glenn Foundation (T.A.), American Federation for Aging Research (T.A.), Hillblom Foundation (G.B.), JSPS (Y.F.), Japanese Biochemistry Postdoctoral Fellowship (Y.F), Japan Agency for Medical Research and Development (AMED) (JP25wm0625523h0001) (T.M.), National Science Foundation (J.S.), Bakar Aging Research Institute (S.A.V.), gift from Marc and Lynne Benioff, and National Institute on Aging (AG081038 (G.B.), AG086042 (J.S.), AG062357 (S.A.V.), AG077816 (S.A.V). We thank the Genomics CoLabs at the UCSF Institute for Human Genetics for assistance and the UCSF CAT core for sequencing. Some computing was performed on the Sherlock cluster, and we thank Stanford University for related computational resources. We thank the Mass Spectrometry Resource at UCSF (A.L.B.), which is supported by the Dr. Miriam and Sheldon G. Adelson Medical Research Foundation and the UCSF Program for Breakthrough Biomedical Research.

## AUTHOR CONTRIBUTIONS

A.P. and S.A.V. developed concept and designed experiments. A.P. collected and analyzed data. A.P. performed molecular, cellular biochemical, *in vivo* studies and behavioral analyses. L.R. assisted with cellular and biochemical analysis. J.M. and A.L.B. performed mass spectrometry analysis. K.J.B.P. assisted with snRNAseq analysis. S.S., T.A., G.B., R.C., G.A., Y.F., R.M and S.M. assisted with molecular, biochemical and behavioral analysis. J.S and J.C. assisted with sequencing analysis. B.Z and X.S.X. performed electrophysiology analysis. S.F. assisted with experimental design. T.M. performed human analysis. A.P. and S.A.V wrote manuscript and supervised all aspects of this project. All authors had the opportunity to discuss results and comment on manuscript.

## SUPPLEMANTARY FIGURE LEGENDS

**Supplementary Figure 1: Single nuclei RNA sequencing of hippocampus and hippocampal oligodendrocytes from young and aged mice. A)** UMAP projection of hippocampal nuclei colored by age. **(B-D)** Violin plots showing the number of unique RNA features (B), unique RNA counts (C), and percentage of mitochondrial RNA (D) per cluster. (E) Dot plot showing representative marker genes for each cell type identified by single-nucleus RNA sequencing of young and aged hippocampus. **F)** qPCR analysis of cell markers for CNS cell types in hippocampal O4^+^ cells, normalized to unsorted hippocampus.

**Supplementary Figure 2: Matrisome gene detection in bulk sequencing and mass-spectrometry datasets. A)** Number of Matrisome genes detected per category in bulk sequencing of O4^+^ sorted cells. **B)** Number of Matrisome proteins detected per category in mass-spectrometry of soluble hippocampal proteins. **C)** Number of Matrisome genes detected per category in hippocampal bulk sequencing of Hapln2 shRNA and scramble mice.

**Supplementary Figure 3: Hapln2 expression analysis with age in male and female mice. A)** Immunofluorescence of HAPLN2 at the node of Ranvier, delineated by CASPR (paranode) and MBP (myelin sheath). **B)** Expression of *Hapln2* from snRNA-seq of young and aged hippocampus. **C)** qPCR analysis of *Hapln2* in the hippocampal tissue of young and aged mice. **D)** Western Blot of HAPLN2 in hippocampal lysates from young and aged female mice. **E)** Quantification of HAPLN2 protein levels shown in (D).

**Supplementary Figure 4: Validation of lentiviral-mediated Hapln2 overexpression in oligodendrocytes of young mice. A)** Schematic illustrating experimental groups. Young mice (3 months) were injected with lentivirus to selectively increase the expression of Hapln2 in oligodendrocytes. **B)** Representative image showing expression of the control lentivirus encoding the recombinant protein mCherry. **C)** qPCR analysis of Hapln2 expression in the hippocampus of mice injected with control or Hapln2 overexpressing lentivirus. N= 12-14.

**Supplementary Figure 5: Oligodendrocyte-specific Hapln2 overexpression in young mice does not affect general health metrics. A)** Schematic illustrating the experimental paradigm of oligodendrocyte-specific Hapln2 overexpression in young mice, followed by subsequent testing of health metrics. **B-C)** Locomotor activity (B), and anxiety-like behaviour (C, time in center) were assessed using open field (OF) assay, during which mice explored an empty arena for 10 min. **D)** Nest forming performance, a hippocampal-dependent well-being metric was assessed using a pre-defined scoring system. **E)** Body weigh measurements. N= 12-14 mice. Data shown as mean +/- s.e.m.

**Supplementary Figure 6: Validation of AAV-mediated Hapln2 knockdown in aged mice. A)** qPCR analysis of Hapln2 expression of HEK cells overexpressing Hapln2 and treated with scramble of Hapln2 shRNA. **B)** Schematic illustrating experimental groups. Aged mice were injected retro-orbitally with scramble or Hapln2 shRNA AAV. **C)** Representative image showing expression of the control AAV encoding the scramble AAV with mCherry. **D)** Western Blot of HAPLN2 in hippocampal lysates of aged mice injected with scramble of Hapln2 shRNA AAV. **E)** Quantification of HAPLN2 protein levels shown in (D).

**Supplementary Figure 7: Hapln2 knockdown in aged mice does not alter inflammatory markers. A)** Representative IF images of IBA-1 and C1q in the hippocampus of aged mice injected with scramble or Hapln2 shRNA AAV. **B)** Quantification of IBA-1 and C1q immunoreactivity.

**Supplementary Figure 8: Single nuclei RNA sequencing of the hippocampus of aged mice following Hapln2 abrogation. A)** UMAP projection of hippocampal nuclei colored by condition. **B-D)** Violin plots showing the number of unique RNA features (C), unique RNA counts (D), and percentage of mitochondrial RNA (E) per cluster. **E)** Dot plot showing representative marker genes for each cell type identified by single-nucleus RNA sequencing of Hapln2 shRNA and scramble mice.

**Supplementary Figure 9: Hapln2 knockdown in aged mice does not affect general health metrics. A)** Schematic illustrating the experimental paradigm of Hapln2 knockdown in aged mice, followed by subsequent testing if health metrics. **B-C)** Locomotor activity (B), and anxiety-like behavior (C, time in center) were assessed using open field (OF) assay, during which mice explored an empty arena for 10 min. **D)** Nest forming performance, a hippocampal-dependent well-being metric was assessed using a pre-defined scoring system. **E)** Body weigh measurements. N= 9-11 mice. Data shown as mean +/- s.e.m.

## METHODS

### Mouse tissue collection

Mice were deeply anesthetized with ketamine (87.5 mg kg⁻¹) and xylazine (12.5 mg kg⁻¹) before transcardial perfusion with ice-cold phosphate-buffered saline (PBS). Brains were rapidly removed, sectioned in half along the sagittal plane, and processed depending on downstream applications. For molecular analyses, hippocampi were sub-dissected on ice and snap-frozen. For histological studies, half brains were immersion-fixed in 4% paraformaldehyde (PFA) in PBS (pH 7.4) at 4 °C for 48 h, followed by cryoprotection in 30% sucrose until equilibration.

### Human tissue sample collection

CNS tissue was obtained from the archives of the Institute of Neuropathology at Kyoto University. This study was approved by the local ethics committee (R1038). Hippocampal samples were obtained from nine autopsy cases, comprising four young (34–45 years) and five aged (89–95 years) individuals without neurological disease.

### Single-nuclei RNA sequencing and analysis

Nuclei were isolated based on the protocol established by 10x Genomics with modifications and performed on nuclei isolated from four-six mice. Two mice were pooled together per group. Briefly, flash frozen dissected hippocampi were dounce homogenized (Wheaton, Cat# 357538) in 500μL of Nuclei EZ prep lysis buffer (Sigma-Aldrich, NUC101, 1x RNAse Inhibitor) with 20 strokes of the loose pestle and 20 strokes of the tight pestle. 500 μL of NP40 lysis buffer was added and samples incubated for 7 minutes on ice. Samples were filtered through a 40 μm filter and centrifuged at 500 RCF for 5 min at 4°C. Samples were resuspended in 1ml of Wash Buffer (PBS, 1%BSA, 1xRNAse Inhibitor) and incubated for 5 min on ice. Samples were centrifuged at 500 RCF for 5 min at 4°C, supernatant was removed, and samples were resuspended in 400 uL of Wash Buffer with 1:10,000 dilution of Hoescht 33342 and incubated for five minutes before filtering through a 35μm FACS tube filter and sorting. Nuclei were sorted on a BD FACSAria II with a 100 μm nozzle and with a flow rate of 1–2.5. Nuclei were first gated by forward and side scatter, then gated for doublets with height and width. Nuclei that were Hoechst+ were sorted and samples were combined per group. Isolated nuclei were given to the UCSF-CoLab Genomics Core for analysis with the 10x Genomics Chromium Single Cell Expression Solution 3’ kit. The Genomics Core prepared cells for 10x Genomics Chromium single-cell capture. 30,000 cells were loaded per sample. cDNA libraries were prepared according to the standard 10x Genomics protocols. The final library pool was sequenced on the NovaSeq600 or the NovaSeqX 10B system at the UCSF CAT Core. The raw base sequence calls were demultiplexed into sample-specific cDNA files with bcl2fastq/mkfastq and converted to count matrices using Cell Ranger 7.1 (10x Genomics) and aligned to the mm10 reference genome. Downstream analysis, including quality control, normalization, dimensionality reduction, clustering, and differential gene expression, were performed in Seurat (v4.3) using standard workflows. Nuclei expressing <300 genes, >6000 genes, or > 10% mitochondrial genes were excluded from analysis. Differential gene expression was calculated using Seurat (min.pct = 0.01, log2 fold change >0.15, pseudocount = 0.1). For comparisons of differentially expressed genes between clusters, cell numbers were downsampled to match the smallest cluster before identifying differentially expressed genes.

### Hippocampal oligodendrocyte cell isolation

Hippocampal oligodendrocytes were isolated from 3-month and 22-month old C57/B6 mice which were decapitated after a lethal injection with ketamine (87.5 mg kg⁻¹) and xylazine (12.5 mg kg⁻¹). We used a previously described method for isolating oligodendrocyte progenitor cells^52^ with some modifications. The hippocampi were quickly dissected on ice, and placed into ice-cold Hibernate-A (Gibco). The hippocampi of 5 mice were pooled together for 1 biological replicate. The hippocampi were mechanically minced into approximately 1mm^3^ pieces on ice. The tissue was collected and spun down at 100xg for 1min at RT and washed in HBSS^-/-^ (no Mg2+ and Ca2+, Gibco). Hippocampi were dissociated using the papin dissociated system (Worthington, cat. no. LK003153). The tissue was dissociated using a shaker (50 rpm) for 40mins at 35°C. The tissue was centrifuged at 200xg for 3 mins at RT, and the supernatant was discarded. Tissue was resuspended in titration solution (Hibernate-A, Pyruvate 2mM, B27 2%, pH 7.3-7.4). To obtain single cell suspension, the tissue was triturated 10 times using a 5mL fire polished glass pippete. The digested supernatant was collected and filtered through a 70 μm cell strainer into tubes that contained a 90% percoll solution (diluted in 10x PBS). The solution was mixed to give a homogenous suspension and adjusted to a final percoll concentration of 22.5% with DMEMF12 (Gibco xxxx). The single cell suspension was then centrifuged at 800xg for 20mins at RT, with no break, to separate remaining debris. All layers without cells were discarded and the cell pellet was resuspended in red blood cell lysis buffer, incubated for 90 seconds at RT to remove red blood cells. 9 mL of HBSS^+/+^ (Gibco) was added to the suspension to detain the reaction, and spun down at 300xg for 5 min at RT. The cell pellet was then resuspended in 0.5mL of wash buffer (2mM EDTA, 2mM Na-Pyruvate, 0.5% BSA in PBS, pH 7.3) and incubated with 2.5uL of anti-O4 microbeads (Miltenyi Biotec). The same was gently mixed and placed in the dark at 4°C for 15 minutes. Cells were washed with 7mL washing buffer and centrifuged at 300xg for 5mins. The cell pellet was resuspended in 0.5mL and MACS was performed according to the manufacturer’s instructions. Briefly, a MS column (Milteny, 130-042-201) was pre-wet with 0.5mL of washing buffer and resuspended cells were transferred to one MS column. The column was then washed with 0.5mL wash buffer for a total of 3 times. Finally, O4+ cells were flushed out of the column with 1mL wash buffer. Subsequently cells were centrifuged (300xg for 5mins at RT) and resuspended in Trizol for subsequent analysis.

### Bulk RNA-seq analysis

Total RNA was extracted from frozen hippocampal tissue using TRIzol reagent (Thermo Fisher) according to the manufacturer’s protocol, followed by chloroform phase separation and isopropanol precipitation. The RNA pellet was washed with 75% ethanol, air-dried, and resuspended in RNase-free water. RNA purity and concentration were assessed by spectrophotometry (Nanodrop one, ThermoScientific). RNA samples were submitted to Azenta Life Sciences (formerly Genewiz, South Plainfield, New Jersey) where RNA quality was confirmed using an Agilent 2100 Bioanalyzer (RIN ≥ 8.0). Library preparation and sequencing were performed using standard Illumina protocols. Reads were aligned to the *mm10* mouse transcriptome using STAR version 2.7.3a^53^ with ENCODE-recommended settings. Gene-level quantification was performed with RSEM version 1.3.1^54^, and differential expression analysis was carried out in R version 4.0.2 using DESeq2 version 1.28.1^55^. The full pipeline details (version 2.1.2) are available at https://github.com/emc2cube/Bioinformatics/. Gene Ontology enrichment was performed using DAVID, with all detected genes in the dataset serving as the background reference list. Matrisome analysis was performed as described in^21^, full pipeline details are available at https://github.com/Matrisome/MatrisomeAnalyzeR.

### Mass Spectrometry

For mass spectrometry analysis, samples originate from^56^ . Mouse hippocampi were isolated from fresh mouse brain tissue of either 3- or 24-month-old mice after perfusion with cold PBS (n= 3 replicated per group, n= 3 mice per replicate). Protein was isolated from the hippocampus with TRIzol Reagent (Thermo Fisher Scientific), according to manufacturer’s instructions. Frozen protein pellets were resuspended in 50mM ammonium bicarbonate containing 6M guanidine hydrochloride, 6X Phosphatase Inhibitor Cocktails II and III (Sigma-Aldrich), and 80mM PUGNAc (Tocris Bioscience, Avonmouth, UK). Protein concentrations were estimated with bicinchoninic acid (BCA) protein assay (ThermoFisher Scientific, Rockford, IL). The protein lysate (1 mg from each sample) was reduced for 1 hour at 56°C with 2.5 mM Tris(2-carboxyethyl)phosphine hydrochloride and subsequently carbamidomethylated using 5 mM iodoacetamide for 45 min at room temperature in the dark. Lysates were diluted to 1M guanidine hydrochloride with 50 mM ammonium bicarbonate, pH 8.0, and equal amounts of each sample were digested overnight at 37°C with sequencing grade trypsin (ThermoFisher Scientific) at an enzyme to substrate ratio of 1:50 (w/w). Following digestion, samples were acidified with formic acid (FA) (Sigma-Aldrich), desalted using a 360-mg C18 Sep-Pak SPE cartridge (Waters), and dried to completeness using a SpeedVac concentrator (Thermo Electron).

For TMT labelling, tryptic peptides were labeled with TMT-10plex according to the manufacturer’s protocol. The TMT labeling was as follows: 3month-1:TMT127C, 3month-2:TMT127N, 3month-3:TMT128C, 12month-1:TMT128N, 12month-2:TMT129C, 12month-3:TMT129N, 24month-1:TMT130C, 24month-2:TMT130N, 24month-3: TMT-131C. Labeling efficiency was checked on Thermo Scientific Q Exactive Plus Orbitrap. TMT labeled peptides were mixed, desalted using a 360-mg C18 Sep-Pak SPE cartridge, and dried to completeness using a SpeedVac concentrator before peptide separation.

Peptides were separated by high pH RPLC using a 150 × 4.6 mm Gemini 5μ C18 column (Phenomenex, Torrance, CA). Peptides were loaded onto the column in 20 mM NH4OCH3, pH 10 (buffer A) and subjected to a gradient from 1% to 9% 20mM NH4OCH3, pH10 in 90% acetonitrile (buffer B) over 2.0 mL, up to 49% B over 20.0 mL, and up to 70% B over 1.5 ml at a flow rate of 0.55 mL/min. Peptides were collected over 48 fractions.

For mass spectrometry analysis, peptide fractions from the high pH RPLC separation were concatenated into 24 fractions and 12 were analyzed on an Orbitrap Fusion Lumos (Thermo Scientific, San Jose, CA) equipped with a NanoAcquity UPLC (Waters, Milford, MA). Peptides were fractionated on a 50cm x 75 μm ID 2μm C18 EASY-Spray column using a linear gradient from 3.5-30% solvent B over 185 min. Precursor ions were measured from 375 to 1500 m/z in the Orbitrap analyzer (resolution: 120,000; AGC: 4.0e5). Each precursor ion (charged 2-7+) was isolated in the quadrupole (selection window: 1.0 m/z; dynamic exclusion window: 30 s; MIPS Peptide filter enabled) and underwent two ms2 fragmentation methods. Ions were fragmented by HCD (Maximum Injection Time: 86 ms; HCD Collision Energy: 35%, Stepped Collision Energy: 5%) and measured in the Orbitrap (resolution: 50,000; AGC; 5.0e4). The scan cycle was 3 seconds.

Peaklists were extracted using Proteome Discoverer 2.2. Data was searched against the SwissProt Mus musculus database (downloaded September 6, 2016) (and concatenated with a randomized sequence for each entry) using Protein Prospector (v5.23.0). Cleavage specificity was set as tryptic, allowing for 2 missed cleavages. Carbamidomethylation of Cys and TMT10plex on lysine and the peptide N-terminus were set as constant modifications. The required mass accuracy was 10 ppm for precursor ions and 30 ppm for fragment ions. Variable modifications include Acetyl (Protein N-term), Acetyl+Oxidation (Protein N-term M), Gln->pyro-Glu (N-term Q), Met-loss (Protein N-term M), Met-loss+Acetyl (Protein N-term M), Oxidation (M), Pyro-carbamidomethyl (N-term C). Two modifications per peptide were permitted.

For TMT mass spectrometry data analysis, data was filtered to only include peptides unique to a single protein. Quantitation of TMT data was performed by calculating ratios of reporter ion peak intensities between conditions along with variance for each ratio, and median normalized. Peptide abundances were normalized by the median of ratio distributions. Age-dependent changes were quantified as the normalized median log₂ fold change between 24-month-old and 3-month-old samples (12-month-old samples were excluded from this analysis). Proteins were considered differentially expressed at *p* < 0.05. Gene Ontology enrichment analysis of differentially expressed proteins was performed using DAVID, with all detected proteins in the dataset serving as the background reference list. Raw data files available under MassIVE submission.

### Viral Production

Adherent HEK293T cells (ATCC) were cultured in Dulbecco’s Modified Eagle Medium (DMEM) supplemented with 10% fetal bovine serum (FBS) and 1% penicillin–streptomycin. Cells were maintained in 10-cm dishes or T175 flasks under standard conditions (37 °C, 5% CO₂). Transfections were performed using Lipofectamine (ThermoFisher) for lentivirus production and polyethylenimine (1 mg mL⁻¹; Polysciences, Cat# 23966-1) for AAV production.

### Lentivirus production

HEK293T cells were transfected with a 4:3:1 µg ratio of lentiviral transfer construct, psPAX2 packaging plasmid (Addgene #12260; gift from D. Trono), and pCMV-VSV-G envelope plasmid (Addgene #8454, RRID:Addgene_8454)^57^. After 24 h, media were replaced with fresh growth medium supplemented with viral boost. Supernatants were collected at 48 h post-transfection and clarified by low-speed centrifugation to remove cell debris.

### AAV production

For AAV generation, HEK293T cells were transfected with the AAV transfer plasmid, PHP.eB capsid plasmid (Addgene #103005; gift from Viviana Gradinaru) ^58^, and helper plasmid using PEI in Opti-MEM (Thermo Fisher Scientific Cat# 31985-062). After overnight incubation, media were replaced with fresh DMEM, and both cells and supernatant were harvested 72 h later for viral isolation. Cells were mechanically detached and combined with the viral-containing media, which were centrifuged for 10 min at 100 × g at 4 °C to remove debris. The remaining cell pellet underwent four freeze–thaw cycles to release intracellular AAV particles, and the lysate was clarified by centrifugation at 7500 x g for 5 mins. The resulting supernatant was recombined with the conditioned media and filtered through a 0.45 µm filter.

### Viral purification and titration

Both lentiviral and AAV preparations were purified and concentrated by ultracentrifugation (Beckman Coulter) at 24,000 rpm for 90 min. Viral pellets were resuspended in sterile PBS. Lentiviral genomic titers were determined using Lenti-X GoStix Plus (TakaraBio), whereas AAV genomic titers were quantified by qPCR using primer sets targeting the *mCherry* sequence (FWD: GACTACTTGAAGCTGTCCTTCC, REV: CGCAGCTTCACCTTGTAGAT). AAV vectors were administered via retro-orbital injection.

### Stereotaxic injections

Procedures were adapted from Lin et al (get reference from Laura’s paper). Mice were anesthetized with 2% isoflurane in oxygen (2 L min⁻¹) and positioned in a stereotaxic frame. Ophthalmic ointment was applied to prevent corneal drying, and fur over the scalp was trimmed. 4 μL of lentiviral solution (1.0 × 10^9^ IFU/mL) was bilaterally injected at the following coordinates relative to bregma: anteroposterior (AP) −2.0 mm, mediolateral (ML) ±1.5 mm, and dorsoventral (DV) −1.8 mm and −2.1 mm from the skull surface. A total volume of 2 µl per site was infused using a 5-µl, 26-gauge Hamilton syringe at a rate of 0.2 µl min⁻¹ over 10 minutes. To prevent reflux along the needle track, the syringe was left in place for 8 minutes following infusion and then partially withdrawn and held for an additional 2 minutes before complete retraction. The incision was closed using sterile silk sutures and sealed with VetBond. Postoperative care included subcutaneous administration of saline, enrofloxacin, carprofen, and buprenorphine, and animals were closely monitored until full recovery.

### Behavioral Analysis

Animals were group-housed until one week before behavioral testing, after which they were singly housed with environmental enrichment (nestlets and shelters). Health and body weight were monitored throughout the course of all experiments. All behavioral testing was conducted in the Villeda Laboratory behavioral suite at the UCSF Parnassus campus.

### Open field test

Spontaneous locomotor activity and anxiety-related behavior were assessed using the open-field test. Mice were placed in a square arena (40 × 40 cm) and allowed to explore freely for 10 minutes. Total distance traveled and time spent in the center versus periphery of the arena were automatically recorded and analyzed using the MotorMonitor software (Kinder Scientific). The apparatus was cleaned with 0.3% acetic acid between trials to remove olfactory cues.

### Novel Object Recognition

Recognition memory was evaluated using a two-trial novel object recognition task in the same open-field arena. On day 1 (familiarization), mice were allowed to explore two identical objects for 5 minutes. After a 24 h retention interval, one familiar object was replaced with a novel object of similar size and texture, and mice were again allowed to explore for 5 minutes. Exploration time directed toward each object was tracked using Smart Video Tracking Software (Panlab, Harvard Apparatus). Mice that failed to explore both objects during the training phase or that exhibited low total object interaction times during testing were excluded from analysis.

### Radial arm water maze

Spatial learning and memory were evaluated using the 8-arm radial arm water maze (RAWM) paradigm as previously described^59^. In this task, mice were trained to locate a hidden escape platform located at the end of a constant goal arm. The start arm varied pseudorandomly between trials. Entry into an incorrect arm was recorded as an error, and errors were averaged across training blocks consisting of three consecutive trials.

On the training day (day 1), mice completed 12 trials (blocks 1–4) alternating between visible and hidden platform trials. After a 1-hour rest period, learning was assessed in three additional hidden-platform trials (block 5). On the testing day (day 2), mice were evaluated in 9 hidden-platform trials (blocks 6–8) to assess memory retention. Performance was analyzed as the mean number of errors per block. All trials were recorded and scored by experimenters blinded to treatment and condition.

### Health metrics

Nest-building behavior was assessed using the nestlet assay as previously described^60^. Mice were provided with two pressed cotton nestlets and allowed up to 48 hours to construct nests. Nest quality was scored on a 1–5 scale, where a score of 1 indicated no nest construction and a score of 5 indicated a fully enclosed nest, as previously defined.

### Immunofluorescence

Tissue processing of mouse hippocampus and IF were performed on free-floating sections. First, brains were sectioned coronally at 40 µm on a freeze-stage microtome (Leica). Free-floating sections were permeabilized in pre-treatment buffer (0.1% Triton X-100 in TBST) for 45 mins, washed three times in TBST, and blocked in TBST containing 3% normal donkey serum (NDS). Sections were then incubated overnight at 4 °C with primary antibodies diluted in TBST + 3% NDS, including anti-HAPLN2 (1:500, Santa Cruz), anti-CASPR (1:2000, abcam), anti-MBP (1:500, BioRad), anti-C1q (Abcam), anti-IBA1 (Synaptic Systems), anti-mCherry (1:2000, abcam). Fluorescent labeling was visualized using Alexa Fluor–conjugated secondary antibodies (1:500; Invitrogen) in TBST + 3% NDS for 1 h at room temperature. Nuclei were counterstained with Hoechst (Thermo Fisher Scientific, Cat# H3570). Sections were mounted onto Superfrost Plus microscope slides and coverslipped with ProLong Gold antifade reagent.

Images were acquired using a Zeiss LSM 800 or LSM 900 confocal microscope. Labeling intensity and thresholded positive area were quantified from 3–5 hippocampal sections per mouse using FIJI (ImageJ) and Zeiss ZEN image-analysis software.

### Immunohistochemistry

Paraffin-embedded human hippocampal sections (6 μm) were deparaffinized and processed for hematoxylin and eosin or Klüver–Barrera staining using standard procedures. For immunohistochemistry, sections were incubated with mouse anti-HAPLN2 (sc-376797, 1:200; Santa Cruz) and rabbit anti-MBP (PD004, 1:200; MBL) primary antibodies. Signal detection was performed using biotinylated secondary antibodies (414151F; Histofine) followed by peroxidase - conjugated avidin and visualization with diaminobenzidine (415194F; Histofine). For double labeling, diaminobenzidine and Fast Red were combined using an alkaline phosphatase–conjugated secondary antibody (MP-5402; Vector Laboratories).

For quantitative analysis, DAB-stained sections were imaged under identical conditions. Images were captured and converted to 8-bit grayscale and binarized in ImageJ using a fixed threshold. The percentage of positive area (%Area) was measured in CA1, CA2, and CA3 regions of interest (ROIs).

### Western Blot

Hippocampi was lysed in chilled RIPA buffer (Abcam, Cat# ab156034) supplemented with complete protease inhibitor (Sigma-Aldrich, Cat# 4693116001) and phosphatase inhibitor (Thermo Fisher Scientific, Cat# 78420). Tissue was homogenized using a Bead Ruptor Elite and ceramic beads (Omni International, Cat# 19-645-3). Crude lysates were centrifuged at 10,000 × g for 10 min at 4 °C to remove cellular debris, and the clarified supernatant was collected. Protein concentrations were determined using a Pierce BCA protein assay (Thermo Fisher Scientific).

Equal amounts of protein were mixed with 4× NuPAGE LDS loading buffer (Invitrogen, Cat# NP0008), resolved on 4–12% SDS–polyacrylamide gradient gels (Bio-Rad, Cat# 11346-02), and transferred to nitrocellulose or PVDF membranes using the Trans-Blot Turbo Transfer System (Bio-Rad). Membranes were blocked for 1 h at room temperature in 5% nonfat dry milk in TBST and incubated overnight at 4 °C with the following primary antibodies: anti-GAPDH (Abcam, ab8245; 1:5,000), anti-HAPLN2 (1:500, ThermoFisher), or anti-Tubulin β3 (1:5000, BioLegend). After washing, membranes were incubated for 1 h at room temperature with HRP-conjugated secondary antibodies: donkey anti-mouse (Invitrogen, A15999; 1:2,000) or donkey anti-rabbit (GE Healthcare, NA934V; 1:2,000). Protein bands were visualized using ECL reagents (Bio-Rad cat #1705060, or Bio-Rad cat #1705060) and imaged with a ChemiDoc Imaging System (Bio-Rad). Band intensities were quantified using the built-in gel analysis tool in FIJI/ImageJ (v2.0.0) and normalized to loading controls.

### RT–qPCR

To quantify mRNA expression levels, equal amounts of cDNA were synthesized using a High-Capacity cDNA Reverse Transcription Kit (Thermo Fisher Scientific, 4368813) and then mixed with SYBR Fast mix (Kapa Biosystems) and primers. β-actin was amplified as an internal control. RT–qPCR was performed in the CFX384 Real Time System (Bio-Rad). Each sample and primer set was run in triplicate, and relative expression levels were calculated using the 2^−ΔΔ*C*t^ method.

### Electrophysiology Brain slice preparation

Acute hippocampal slices were prepared from 23–24-month-old mice, 1 month after retro-orbital injection of *Hapln2* shRNA or scramble AAV. All procedures were approved by the IACUC of AfaSci (protocol no. 0223). Mice were deeply anesthetized with halothane and decapitated. Brains were rapidly removed and immersed in ice-cold, oxygenated artificial cerebrospinal fluid (ACSF) containing (in mM): NaCl 130, KCl 2.5, KH₂PO₄ 1.2, CaCl₂ 2.4, MgSO₄ 1.3, NaHCO₃ 26, and glucose 10 (pH 7.4). ACSF was continuously bubbled with 95% O₂ and 5% CO₂ throughout all procedures. Transverse hippocampal slices (400 µm for CA1 recordings and 500 µm for CA3 recordings) were prepared using a vibrating tissue slicer (Stoelting Co., IL) and allowed to recover for at least 1 h at room temperature in oxygenated ACSF before recording.

### Field potential recordings and input/output analysis

Slices were transferred to a submerged recording chamber (Harvard Apparatus) perfused with oxygenated ACSF (flow rate: 1.75 ml min⁻¹) at room temperature. Field excitatory postsynaptic potentials (fEPSPs) were recorded using glass microelectrodes (1–3 MΩ) filled with ACSF and connected to an Axopatch-2B amplifier (Axon Instruments). Data were digitized using a Digidata 1320A interface and acquired at 10 kHz with pClamp 10.4 software (Molecular Devices).

Biphasic current pulses (0.2 ms per phase; 0.4 ms total) were delivered every 10 s through a concentric bipolar stimulating electrode (FHC, Inc.) positioned in the CA3 region. The recording electrode was placed ∼100 µm away within the same layer. Input/output (I/O) curves were generated by increasing the stimulus intensity in 23 incremental steps of ∼0.1 µA, starting from threshold level (typically 20–50 µA) until the maximal fEPSP response was reached. Excessive stimulation was avoided to prevent fiber damage. The slope of the fEPSP was measured from the initial linear rising phase of the negative deflection using Clampfit 10.4. Each data point represented the average of three consecutive traces. For each slice, the I/O relationship was plotted as fEPSP slope versus stimulus intensity, and curves were averaged across slices from each mouse. Data were analyzed using Clampfit 10.4.

### Data analysis, statistics, and reproducibility

All experiments were randomized and blinded by an independent researcher. Experimenters remained blinded throughout histological, biochemical, and behavioral assessments, and group identities were revealed only after completion of statistical analyses. Data are presented as mean ± s.e.m. The distribution of each dataset was assessed for normality using either the D’Agostino–Pearson omnibus test or the Shapiro–Wilk test. Statistical analyses were performed using GraphPad Prism (versions 8–10; GraphPad Software).

Comparisons between two groups were conducted using two-tailed unpaired Student’s *t*-tests. Comparisons among multiple groups were analyzed using one-way ANOVA followed by the appropriate post hoc test, as indicated in the figure legends. Additional statistical information, including exact *n* values and significance thresholds, is provided in the corresponding figure legends. All data generated or analyzed during this study are included in this article.

