## Supplementary figures and images for "COUNTERING AGE-ASSOCIATED ALTERATIONS IN OLIGODENDROCYTE-DERIVED EXTRACELLULAR MATRIX REJUVENATES COGNITION"

### Supplementary Figure 1

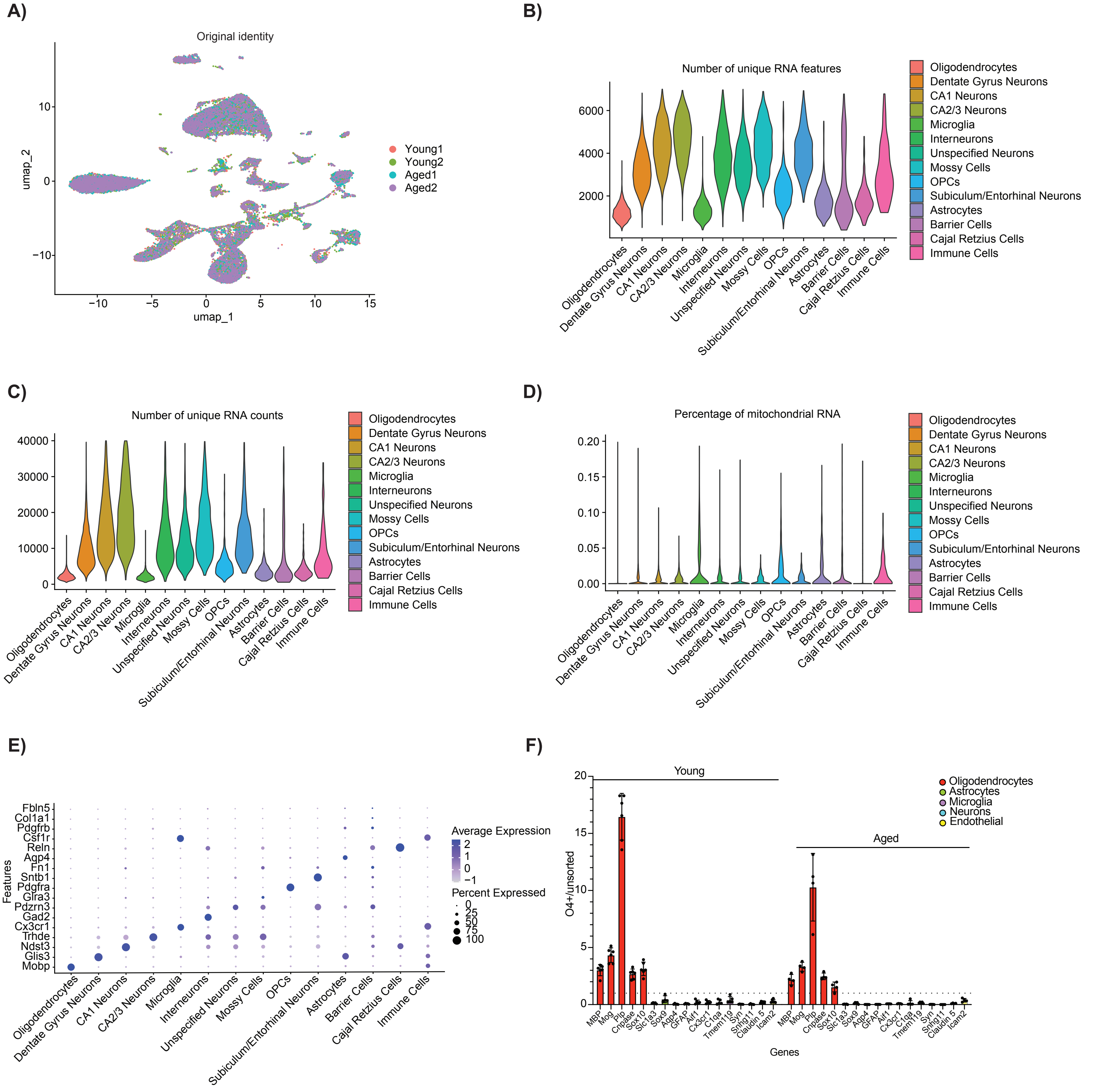


Supplementary Figure 1

### Supplementary Figure 2

A)

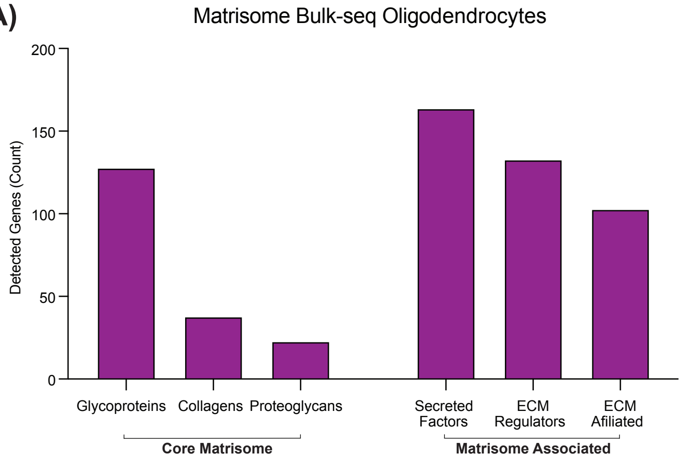


B)

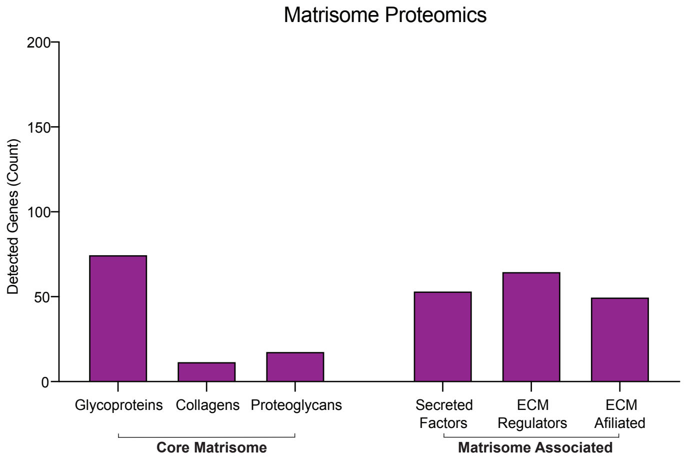


B)

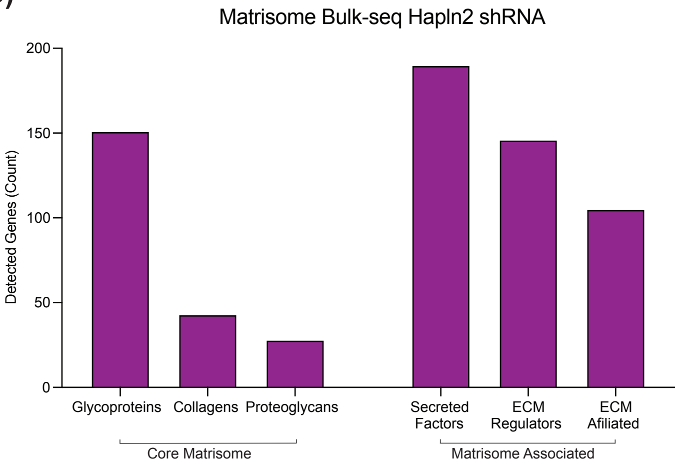


Supplementary Figure 2

### Supplementary Figure 3

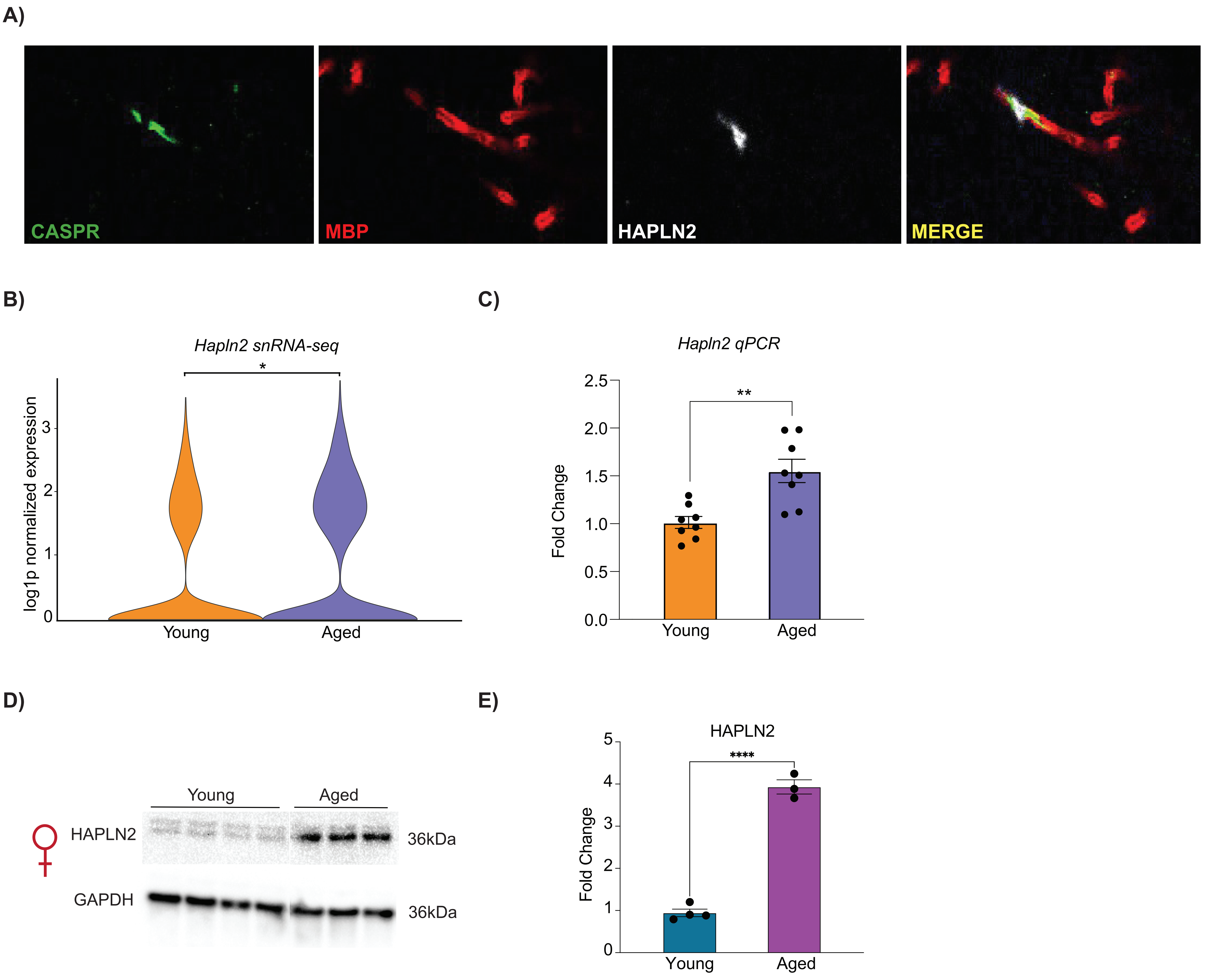


Supplementary Figure 3

### Supplementary Figure 4

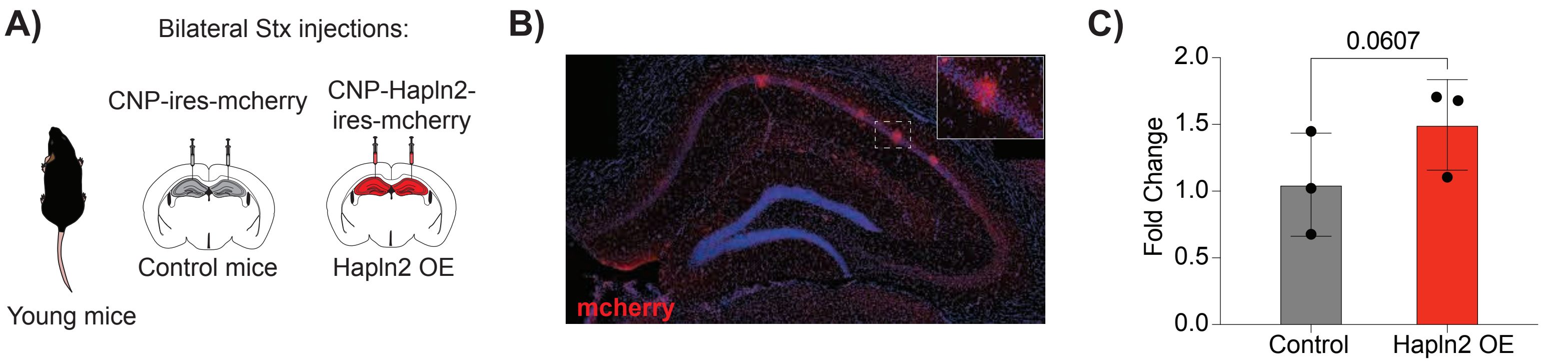


Supplementary Figure 4

### Supplementary Figure 5

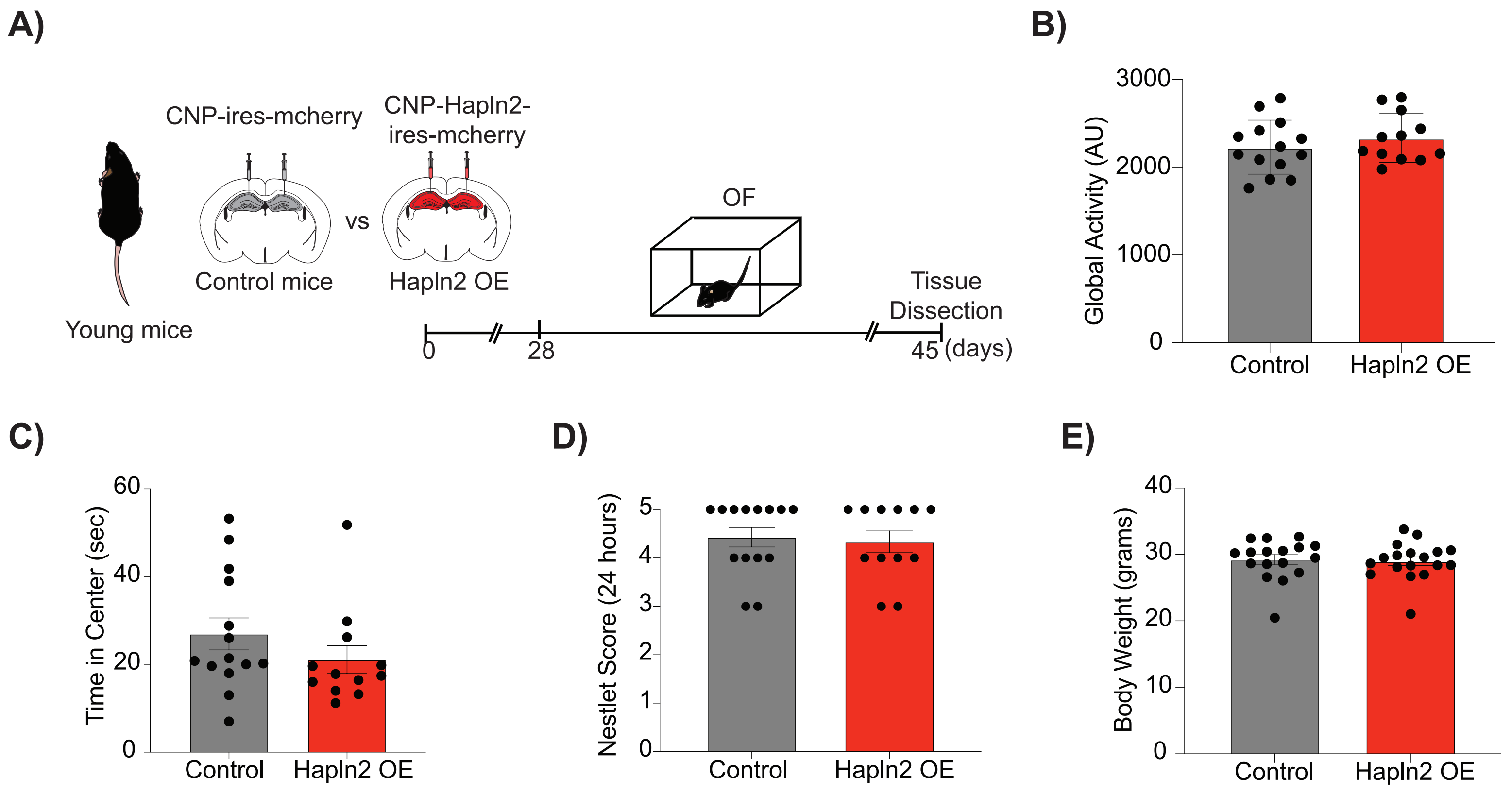


Supplementary Figure 5

### Supplementary Figure 6

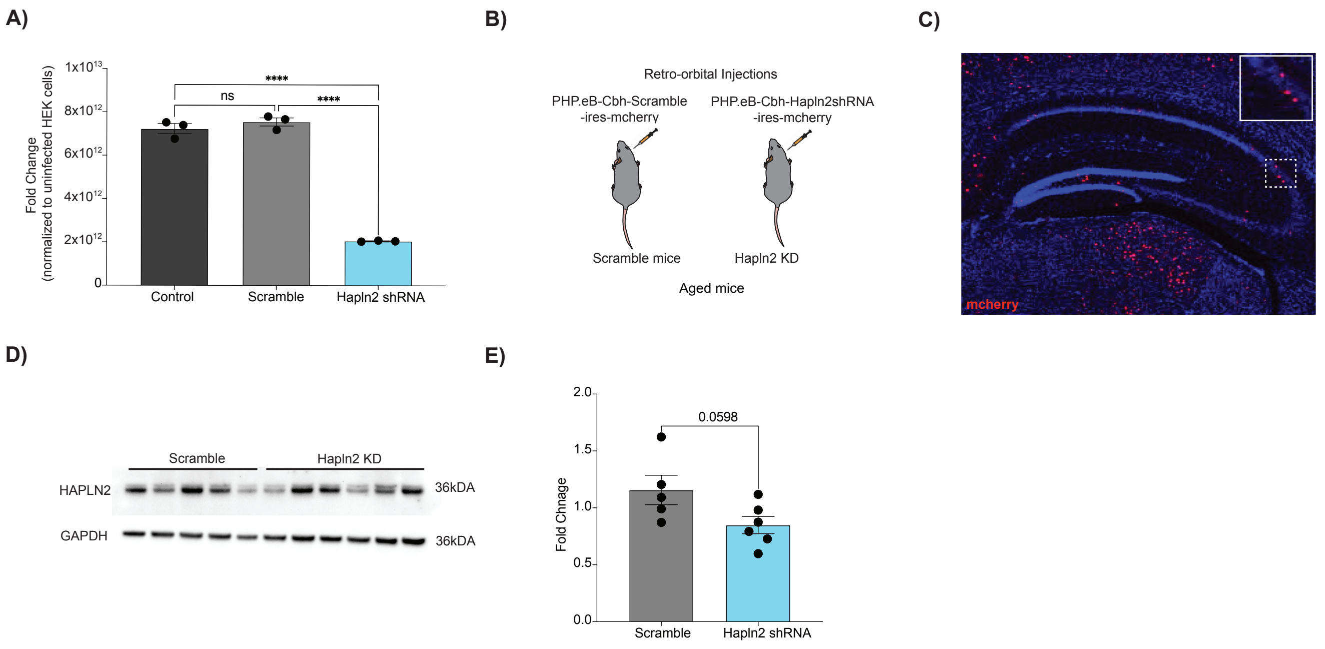


Supplementary Figure 6

### Supplementary Figure 7

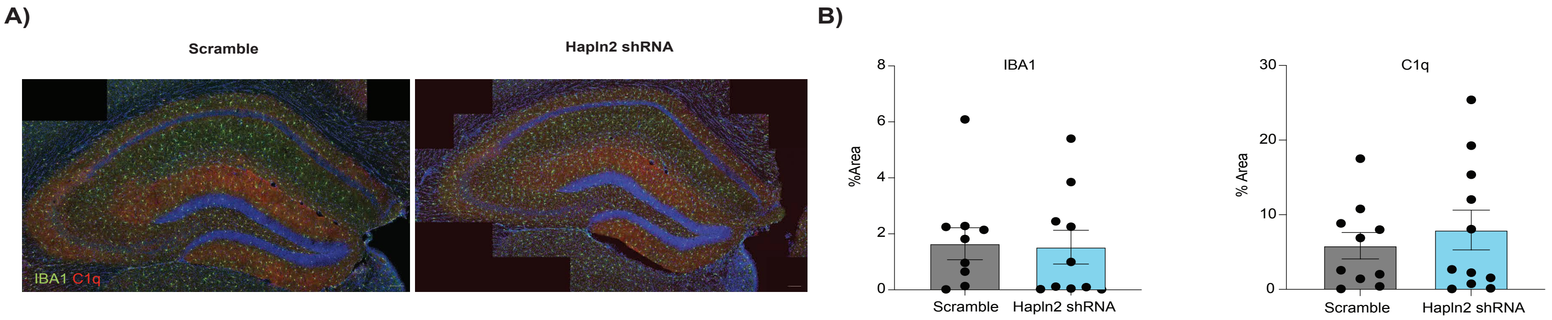


Supplementary Figure 7

### Supplementary Figure 8

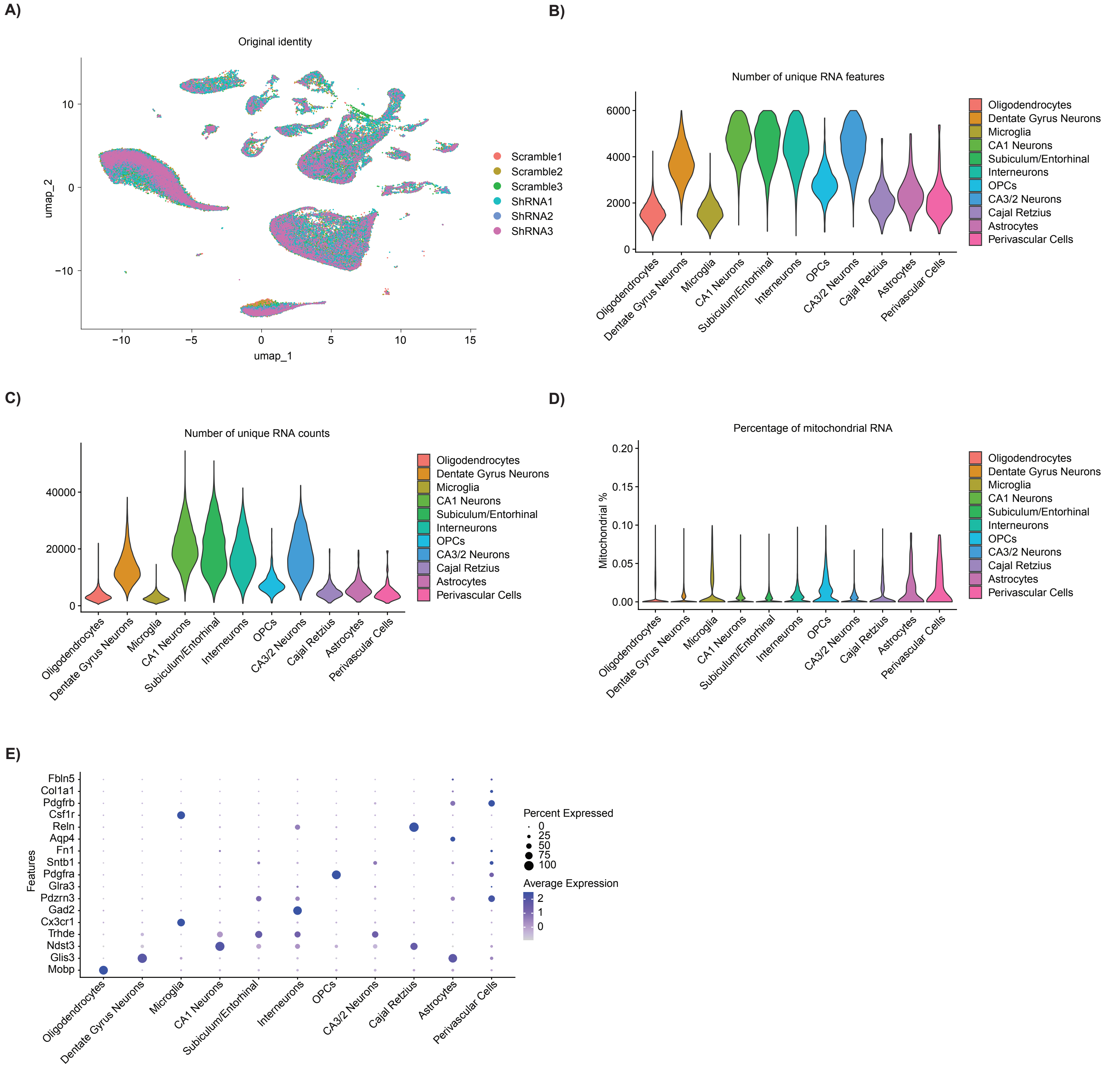


Supplementary Figure 8

### Supplementary Figure 9

A)

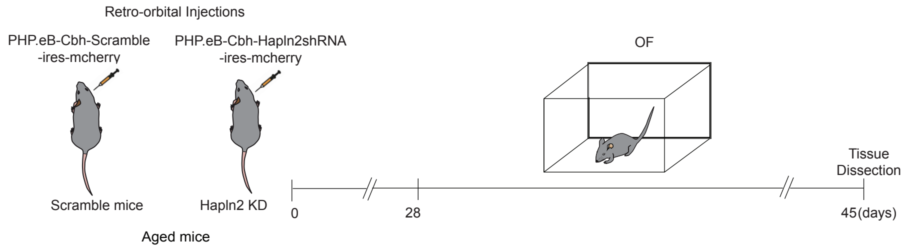


B)

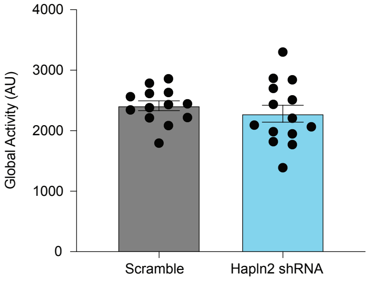


C)

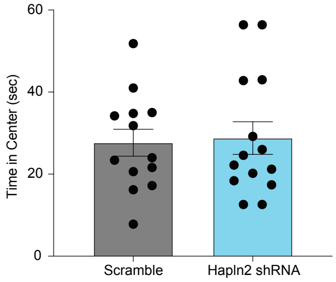


D)

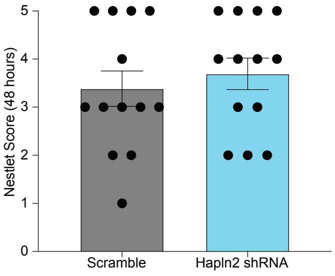


E)

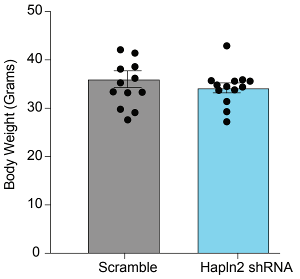


Supplementary Figure 9
